# Negative screening for 12 rare LRRK2 pathogenic variants in a cohort of Nigerians with Parkinson’s disease

**DOI:** 10.1101/2020.06.30.179739

**Authors:** Mie Rizig, Oluwadamilola O. Ojo, Alkyoni Athanasiou-Fragkouli, Osigwe P. Agabi, Olajumoke O. Oshinaike, H Houlden, Njideka U. Okubadejo

**Affiliations:** Department of Molecular Neuroscience, UCL Institute of Neurology and The National Hospital for Neurology and Neurosurgery, Queen Square, London, United Kingdom; Neurology Unit, Department of Medicine, Faculty of Clinical Sciences, College of Medicine, University of Lagos, Lagos State, Nigeria; Neurology Unit, Department of Medicine, Lagos University Teaching Hospital, Idi Araba, Lagos State, Nigeria; Neurology Unit, Department of Medicine, Lagos State University College of Medicine, Ikeja, Lagos State, Nigeria

**Keywords:** Parkinson’s disease, African, black, Nigeria, *LRRK2*, genetic, genes

## Abstract

Mutations in the leucine-rich repeat kinase 2 (*LRRK2*) gene are the most commonly identified genetic variants in familial and sporadic Parkinson’s disease (PD). Over three hundred *LRRK2* variants have been described in the literature, of which at least 17 have a confirmed or probable pathogenic role in PD. The distribution of these rare pathogenic variants has been shown to be different among ethnic groups including Caucasians, Latin Americans and East and South Asians. However, to date no PD-related *LRRK2* pathogenic variant has been described in persons of black African ancestry within or outside Africa. We previously reported that the *LRRK2 p.gly2019ser* mutation was not found in 126 PD patients and 55 controls from Nigeria. Using Kompetitive Allele-Specific Polymerase chain reaction (KASP), we screened a new cohort of 92 Nigerians with PD and 210 healthy ethnically matched controls for 12 rare *LRRK2* variants (which have been shown to be pathogenic in other ethnic populations) including: *p.gly2019ser, p.Arg1441His, p.Gly2385Arg, p.Ala419Val, p.Arg1628Pro, p.Pro755Leu, p.Ile2020Thr* and *Tyr1699Cys*. All 12 rare variants were absent in PD patients and controls from this cohort. These results endorse our previous findings and confirm that rare *LRRK2* pathogenic variants reported in Caucasians, Asians and persons of mixed ancestry are absent in West Africans. Applying next generation sequencing technologies in future studies is necessary to explore possible novel *LRRK2* variants indigenous to black Africans.

## Introduction

The current concept of Parkinson’s disease (PD) is that of a complex disease that results from a combination of genetic and environmental influences. Although PD is the fastest growing neurological disorder and the second most prevalent neurodegenerative disease in Africa, very little is known about the genetic contributors to PD in black Africans. (1, 2) The urgent need to expand the exploration of PD genetics research in Africa is justified by the fact that advances in gene discovery initiatives that occurred in the last twenty years are currently being translated to drug treatments and other preventive and diagnostic interventions. (3) The relevance of these developments to black Africans will depend on understanding the genetic architecture of PD within the continent’s indigenous populations.

Mutations in the leucine-rich repeat kinase 2 (*LRRK2*) gene are the most commonly identified genetic variants in familial and sporadic PD. To date, over three hundred PD-related *LRRK2* variants have been described. (4) The spectrum of these variants ranges from rare mutations with very large effects (i.e. fully or highly penetrant) to genetic variants exerting only modest effects but that are relatively common in the general population. (5) At least 17 rare *LRRK2* variants have demonstrated convincing evidence of pathogenicity in PD based on association, disease segregation and functional studies.(6 – 8) The frequency of these rare variants differs significantly between ethnic groups, suggesting population specific founder effects.(7) For instance, the *p.gly2019ser* - the most commonly reported *LRRK2* PD causing variant worldwide - has it’s highest prevalence in PD in North Africa, where it is reported to occur in 30-41% of familial and 30-39% of apparently sporadic PD respectively. (9) The *p.gly2019ser* contributes to a lesser extent to PD risk in Caucasians (1-4%) and west Asians (1%). (10, 11) The *p.gly2019ser* has not been reported in East Asians and black Africans. (6, 11, 12) Three rare *LRRK2* variants (*p.Gly2385Arg, p.Ala419Val and p.Arg1628Pro*) were repeatedly reported to increase PD risk in East Asians and have been consistently absent in Caucasian PD cohorts. (7) On the other hand, three rare *LRRK2* variants known to occur within the same codon (*p.Arg1441Gly, p.Arg1441Cys* and *p.Arg1441His*) were reported in persons with PD of Caucasian and East Asian origins. (6) Interestingly, the *LRRK2 p.Arg1441Gly* is almost exclusively restricted to the Spanish Basque region with only five reported probands outside Northern Spain. (13 – 18) Similarly, the *p.Ile2020Thr* substitution was only identified in three Japanese pedigrees. (19, 20)

To date no *LRRK2* pathogenic variant has been identified in PD amongst persons of black African ancestry within or outside Africa. A few studies including two from our group in Nigeria, one in Zambia, one in Ghana and several in South Africa have investigated the *LRRK2* PD pathogenic variants in Sub-Saharan Africa. (21–30) In the present report, we expand on our previous enquiry which focused only on *LRRK2 p.gly2019ser* in Nigeria, to explore the frequency of 12 pathogenic rare PD-associated *LRRK2* variants including the *p.gly2019ser* in a new cohort of 92 Nigerians with PD and 210 ethnically matched controls.

## Materials and Methods

This study was approved by the Health Research Ethics Committee of the Lagos University Teaching Hospital, Idi Araba, Lagos State, Nigeria (Ethics approval # ADM/DCST/HREC/366) and the ethics committee of the University College London and the National Hospital for Neurology and Neurosurgery, London, United Kingdom (Ethics approval # 07/Q0512/26). Written informed consent was obtained from all participants.

### Participant recruitment, diagnostic ascertainment and clinical evaluation

Persons with PD were recruited from the Movement Disorders Clinic at the Lagos University Teaching Hospital as part of an ongoing study of the genetics of PD in Nigeria. Case ascertainment was based on the United Kingdom Parkinson Disease Brain Bank (UKPDBB) clinical criteria applied by a Movement Disorders neurologist. (31) Controls were unrelated community volunteers without clinically evident neurological symptoms or signs, otherwise healthy, and matched by gender and local ethnicity. Baseline demographics including age at study, age at onset of PD, gender, family history of PD, and PD stage on the Hoehn and Yahr scale were documented. Clinical characteristics of the cohort are summarized using appropriate descriptors including mean± standard deviation (SD) for continuous variables and % frequency for categorical variables. Statistical analysis was conducted using the Statistical Package for Social Sciences (SPSS)^®^ version 21.0 software.

### Genetic analysis

DNA was extracted from venous whole blood samples (10mls) using standard protocols. Genotyping was performed using Kompetitive Allele-Specific Polymerase chain reaction (PCR) assay (KASP™, LGC Genomics. Herts, UK) as described previously. (22, 32) 12 single nucleotide polymorphisms (SNPs) were selected for genotyping namely: rs34637584 (*p.Gly2019Ser*), rs34995376 (*p.Arg1441His*), rs74163686 (*p.Asn1437His*), rs35870237 (*p.Ile2020Thr*), rs35801418 (*p.Tyr1699Cys*), rs34410987 (*p.Pro755Leu*), rs34778348 (*p.Gly2385Arg*), rs281865052 (*p.Met1869Val*), rs34594498 (*p.Ala419Val*), rs33949390 (*p.Arg1628Pro*), rs35808389 (*p.Leu1114Leu*), rs34805604 (*p.Ile1122Val*). SNPs selection was based on evidence of pathogenicity in PD through replicated association studies, excellent disease segregation data and (or) functional data, in addition to the availability of a validated KASP assay for each SNP. Pathogenic evidence was confirmed by screening ClinVar and MDSGene databases. (33, 34) SNP rs33939927 encoding for *p.Arg1441Gly/Cys/Ser* was originally considered for genotyping but later on deemed unsuitable for the KASP assay due to its close proximity to rs34995376 (*p.Arg1441His*) and it’s tri-allelic nature. Genotyping results were visualized with SNP Viewer software (version 1.99, Hoddesdon, UK). Genotypes were examined for deviation from Hardy-Weinberg equilibrium using Chi-square test.

## Results

92 participants with PD (76.1% male) and 210 controls (66.7% male) were included. All participants were native Nigerians, predominantly of Yoruba ethnicity (62 PD (67.4%) and 172 controls (81.9%)). Mean age at study (years) was 62.1± 9.2 in PD and 64.5 ± 6.7 in controls. 5 (5.4 %) and 20 (21.7%) had early onset PD starting before age 40 and 50 respectively. Positive family history of PD was documented in 9 PD (9.8%) including only 3 with early onset disease. Clinical characteristics of the cohort are summarised in Table 1.

**Table 1.**
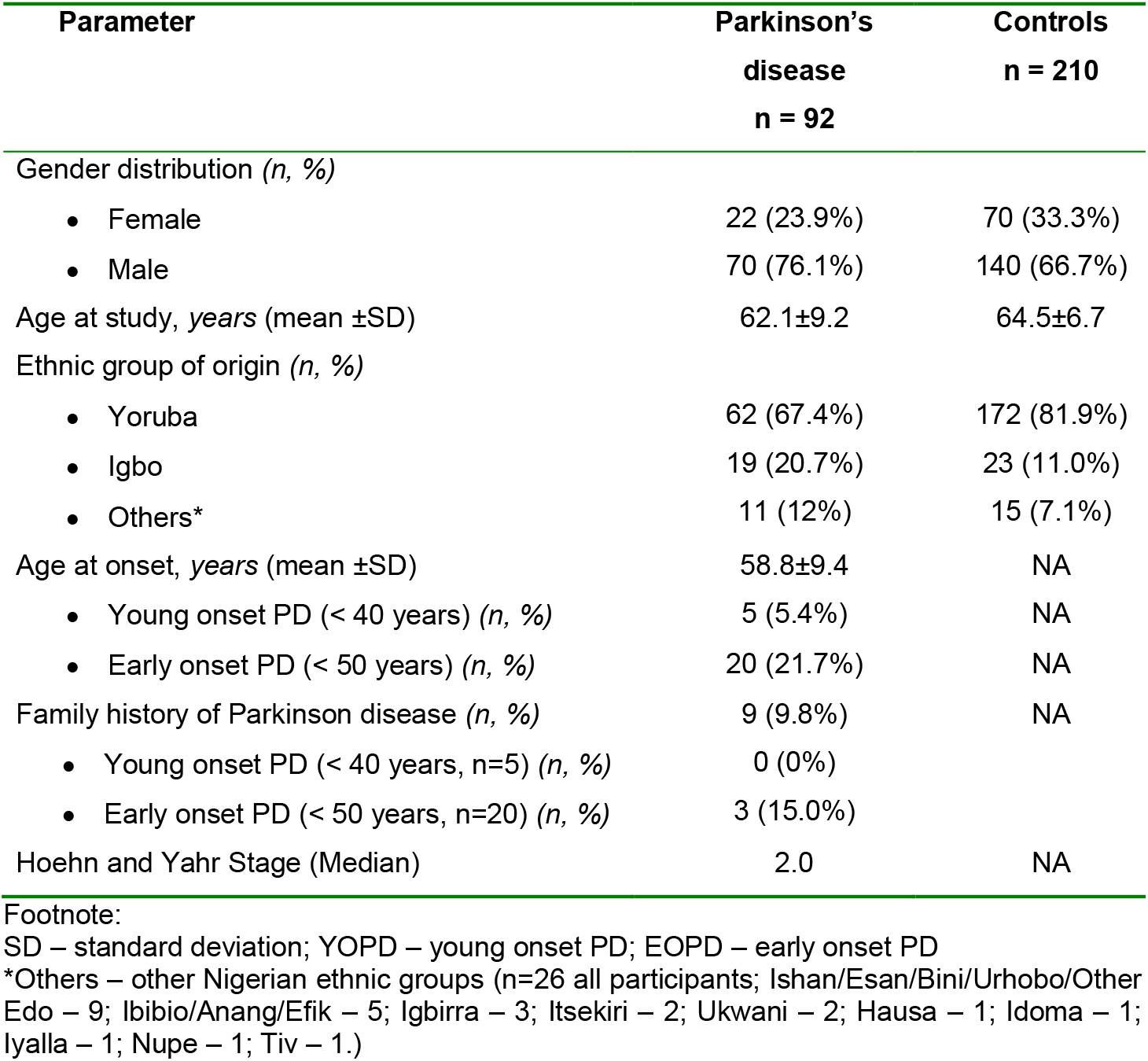
Clinical characteristics of Parkinson’s disease and control participants.

KASP assay performed very well, with 99-100% call rates achieved with clear genotyping clusters and no ambiguous calls for all 12 SNPs. Genotyping calls including uncalled samples from all SNPs are listed in Supplementary Table 1. *LRRK2* pathogenic alleles were absent for all the 12 genotyped SNPS in all PD and control participants. Allelic frequencies for all SNPs were in Hardy-Weinberg equilibrium (P = 1). Details of the 12 SNPs included in this study and their genotyping results are shown in Table 2.

**Table 2.** Description and genotyping results of the 12 *LRRK2* rare SNPs screened in the 92 Nigerian PD patients and 210 controls.

| SNP<br>Allele<br>Position<sup>(1)</sup> | Protein | Type<br>Mutation | Ethnic<br>population at<br>risk of PD | SNP alternative allele frequencies as reported in the Allele Frequency<br>Aggregator (ALFA) <sup>(2)(3)</sup> |  |  |  |  |  | Nigerian<br>cohort |
| --- | --- | --- | --- | --- | --- | --- | --- | --- | --- | --- |
|  |  |  |  | European | African | Asian | Latin<br>American | Others | Total | Genotype |
| rs34594498<br>C>T<br>chr12:40252984 | *p.Ala419Val* | Missense,<br>pathogenic | East Asian | 0.00026 | 0.000 | 0.012 | 0.000 | 0.0007 | 0.00033 | CC<br>(PD 92, Control 210) |
| rs34410987<br>C>T<br>chr12:40283897 | *p.Pro755Leu* | Missense,<br>possibly<br>pathogenic | East Asians | 0.00000 | 0.000 | 0.018 | 0.00 | 0.0003 | 0.00005 | CC<br>(PD 92, Control 210) |
| rs33949390<br>G>A,C,T<br>chr12:40320043 | *p.Arg1628His*<br>*p.Arg1628Pro*<br>*p.Arg1628Leu* | Missense,<br>probably<br>pathogenic | East Asian | A=0.0000,<br>C=0.0005 | A=0.000,<br>C=0.000 | A=0.00,<br>C=0.05 | A=0.000,<br>C=0.000 | A=0.0000,<br>C=0.0000 | A=0.00000,<br>C=0.00063 | GG<br>(PD 92, Control 209) |
| rs35801418<br>A>G<br>chr12:40321114 | *p.Tyr1699Cys* | Missense,<br>pathogenic | European<br>South Korean | 0.00005 | 0.00 | 0.00 | 0.00 | G=0.0000 | 0.00004 | AA<br>(PD 92, Control 210) |
| rs34637584<br>G>A<br>chr12:40340400 | *p.Gly2019Ser* | Missense,<br>pathogenic | European,<br>West Asian and<br>mixed<br>populations | 0.00118 | 0.001 | 0.000 | 0.000 | A=0.0020 | 0.00125 | GG<br>(PD 92, Control 210) |
| rs35870237<br>T>C<br>chr12:40340404 | *p.Ile2020Thr* | Missense,<br>pathogenic | East Asian<br>(Japan/Taiwan) | 0.000 | 0.000 | 0.000 | 0.000 | C=0.0003 | 0.00004 | TT<br>(PD 92, Control 208) |
| rs34778348<br>G>A<br>chr12:40363526 | *p.Gly2385Arg* | Missense,<br>pathogenic | East Asian | 0.000 | 0.000 | 0.038 | 0.003 | 0.0004 | 0.00013 | GG<br>(PD 92, Control 209) |
| rs35808389<br>A>G<br>chr12:40298488 | *p.Leu1114Leu* | Synonymous<br>pathogenic | Europeans | 0.00017 | 0.000 | 0.000 | 0.000 | 0.000 | 0.00015 | AA<br>(PD 92, Control 210) |
| rs34805604<br>A>G<br>chr12:40299125 | *p.Ile1122Val* | Missense,<br>pathogenic | European | 0.00005 | 0.000 | 0.000 | 0.000 | 0.000 | 0.00004 | AA<br>(PD 92, Control 208) |
| rs74163686<br>A>C<br>chr12:40309225 | *p.Asn1437His* | Missense,<br>pathogenic | European | 0.000 | 0.000 | 0.000 | 0.000 | 0.000 | 0.000 | AA<br>(PD 92, Control 210) |
| rs34995376<br>G>A<br>chr12:40310435 | *p.Arg1441His* | Missense,<br>pathogenic | European | 0.00004 | 0.000 | 0.000 | 0.000 | 0.000 | 0.00003 | GG<br>(PD 92, Control 210) |
| rs281865052<br>A>G<br>chr12:40323255 | *p.Met1869Val* | Missense,<br>pathogenic | European<br>(Norway) | 0.000 | 0.000 | 0.000 | 0.000 | 0.000 | 0.000 | AA<br>(PD 92, Control 208) |

Footnote: (1) Position is based on GRCh38. (2) *L. Phan, Y. Jin, H. Zhang, W. Qiang, E. Shekhtman, D. Shao, D. Revoe, R. Villamarin, E. Ivanchenko, M. Kimura, Z. Y. Wang, L. Hao, N. Sharopova, M. Bihan, A. Sturcke, M. Lee, N. Popova, W. Wu, C. Bastiani, M. Ward, J. B. Holmes, V. Lyoshin, K. Kaur, E. Moyer, M. Feolo, and B. L. Kattman. "ALFA: Allele Frequency Aggregator." National Center for Biotechnology Information, U.S. National Library of Medicine, 10 Mar. 2020. Available at: www.ncbi.nlm.nih.gov/snp/docs/gsr/alfa/.* (3) Frequencies are correct for June 2020.

## Discussion

In this study we expanded on our previous enquiry regarding the role of rare *LRRK2* variants in PD in persons of sub-Saharan African black ancestry by exploring a third cohort from Nigeria. We used an efficient, economical and accurate method – the Kompetitive Allele-Specific Polymerase chain reaction (PCR) assay (KASP) – to screen a new cohort of 92 PD patients and 210 controls for five rare variants classified as definitely pathogenic in Europeans, Asians and patients of mixed ancestries: *p.Gly2019Ser, p.Arg1441His, p.Asn1437His, p.Ile2020Thr* and *p.Tyr1699Cys*. In addition we screened for variants known to be pathogenic in East Asians: *p.Gly2385Arg, p.Ala419Val and p.Arg1628Pro*. We added four extremely rare variants to the screening: *p.Pro755Leu, p.Leu1114Leu, p.Ile1122Val* and *p.Met1869Val*. These variants are classified as pathogenic and (or) probably pathogenic. All 12 rare variants were found to be absent in all participants.

The last two decades have witnessed genetic discoveries which have transformed our understanding of PD pathology and pathogenesis. Mutations in the *LRRK2* gene are among the most important findings in PD genetics. Pathogenic mutations in *LRRK2* enhance the enzymatic activity of the *LRRK2* protein, and drugs which inhibit or regulate this activity could have a neuroprotective effect in PD. (6) The genetic contribution of the *LRRK2* to PD pathogenesis in Sub-Saharan Africa remains unknown, and evidence from familial and case control studies in genetically diverse populations demonstrates that *LRRK2* pathogenic variants - with the exception of the *p.Gly2019Ser* - are ethnically or geographically specific. (7)

In the last 12 years a handful of studies screened for *LRRK2* variants in PD cohorts from Nigeria, Zambia, South Africa and Ghana. Yonova-Doing et al used Sanger sequencing to screen for pathogenic variants in *LRRK2* exons 29 to 48 in thirty-eight Zambian patients with idiopathic PD, one patient with juvenile-onset levodopa-responsive atypical Parkinsonism, and 181 ethnically matched controls. (28) The whole *LRRK2* coding region (51 exons) was sequenced in the familial patients and those with onset <55 years (n = 22). No pathogenic variant was identified in any of the Zambian patients and controls although one novel missense variant with unconfirmed pathogenicity (*p.Ala1464Gly*) was reported in one person with apparently idiopathic PD. Cilia et al screened exons 31 (including *p.Gly2019Ser* and *p.Ile2020Thr*) and exon 41 (including, *p.Arg1441gly/Cys/His*) in 54 PD and 46 controls from Ghana, but no pathogenic variant was identified. (29) Data from the studies from South Africa collectively included about 177 PD patients of black South African origin, 14 Nigerians and no controls, although it is unclear if some of the patients were investigated more than once. A combination of methods including multiplex ligation-dependent probe amplification (MLPA), high-resolution melt (HRM), quantitative real time PCR and targeted sequencing panels were used. In all 191 black African ancestry patients, no *LRRK2* pathogenic variant was identified. (29) In 2008, our group applied Sanger sequencing to screen for pathogenic mutations in exons 31 and 41 of the *LRRK2* gene in a cohort of 57 Nigerians with PD and 51 matched controls, and in 2018 we further screened for the *p.Gly2019Ser* in a second cohort of 126 Nigerian PD patients and 55 controls. (21, 22) We were unable to identify any pathogenic variants in both cohorts. In total, our group has screened 275 Nigerians with PD and 316 controls, and no pathogenic variants of LRRK2 have as yet been demonstrated.

The main limitation of this study is the use of a targeted screening methodology (KASP assay) which screened for 12 variants but could not exclude other possible rare mutations such as *p.Arg1441Gly/Cys/Ser, p.Glu923His, p.Lys616Arg, p.Arg151Glu* or identify new pathological variants specific to the cohort. We acknowledge the modest size of the cohort and the importance of explorations including larger samples.

In conclusion, our study corroborates previous reports from our group and others in Sub-Saharan Africa and recaps that rare *LRRK2* variants reported in PD from other populations are as yet unidentified in black Africans. As recently reiterated, only a fraction of likely genetic risk for PD (including that attributable to *LRRK2*) has been identified, and conducting studies in diverse populations such as persons with black African ancestry may hold the key to further unravelling discoveries that will enhance the translation and application of the knowledge into therapies for PD. (35) In order to achieve this and understand the role of *LRRK2* in PD in Africa, resources need to be committed to build sufficiently powered large cohorts and employ advanced sequencing technologies to investigate the role of rare LRRK2 and other PD-related genetic native to black Africans.

---

**Supplementary Table 1.** Genotype calls for all 12 LRRK2 SNPs included in the study

## Notes

### Competing Interest Statement

The authors have declared no competing interest.

