## Supplemental Table 1 for "Negative screening for 12 rare LRRK2 pathogenic variants in a cohort of Nigerians with Parkinson’s disease"

| **Sample ID** | **rs34594498** | **rs34410987** | **rs33949390** | **rs35801418** |
|---|---|---|---|---|
| PD_01 | C:C | C:C | G:G | A:A |
| PD_02 | C:C | C:C | G:G | A:A |
| PD_03 | C:C | C:C | G:G | A:A |
| PD_04 | C:C | C:C | G:G | A:A |
| PD_05 | C:C | C:C | G:G | A:A |
| PD_06 | C:C | C:C | G:G | A:A |
| PD_07 | C:C | C:C | G:G | A:A |
| PD_08 | C:C | C:C | G:G | A:A |
| PD_09 | C:C | C:C | G:G | A:A |
| PD_10 | C:C | C:C | G:G | A:A |
| PD_11 | C:C | C:C | G:G | A:A |
| PD_12 | C:C | C:C | G:G | A:A |
| PD_13 | C:C | C:C | G:G | A:A |
| PD_14 | C:C | C:C | G:G | A:A |
| PD_15 | C:C | C:C | G:G | A:A |
| PD_16 | C:C | C:C | G:G | A:A |
| PD_17 | C:C | C:C | G:G | A:A |
| PD_18 | C:C | C:C | G:G | A:A |
| PD_19 | C:C | C:C | G:G | A:A |
| PD_20 | C:C | C:C | G:G | A:A |
| PD_21 | C:C | C:C | G:G | A:A |
| PD_22 | C:C | C:C | G:G | A:A |
| PD_23 | C:C | C:C | G:G | A:A |
| PD_24 | C:C | C:C | G:G | A:A |
| PD_25 | C:C | C:C | G:G | A:A |
| PD_26 | C:C | C:C | G:G | A:A |
| PD_27 | C:C | C:C | G:G | A:A |
| PD_28 | C:C | C:C | G:G | A:A |
| PD_29 | C:C | C:C | G:G | A:A |
| PD_30 | C:C | C:C | G:G | A:A |
| PD_31 | C:C | C:C | G:G | A:A |
| PD_32 | C:C | C:C | G:G | A:A |
| PD_33 | C:C | C:C | G:G | A:A |
| PD_34 | C:C | C:C | G:G | A:A |
| PD_35 | C:C | C:C | G:G | A:A |
| PD_36 | C:C | C:C | G:G | A:A |
| PD_37 | C:C | C:C | G:G | A:A |
| PD_38 | C:C | C:C | G:G | A:A |
| PD_39 | C:C | C:C | G:G | A:A |
| PD_40 | C:C | C:C | G:G | A:A |
| PD_41 | C:C | C:C | G:G | A:A |
| PD_42 | C:C | C:C | G:G | A:A |
| PD_43 | C:C | C:C | G:G | A:A |
| PD_44 | C:C | C:C | G:G | A:A |
| PD_45 | C:C | C:C | G:G | A:A |
| PD_46 | C:C | C:C | G:G | A:A |

| Sample ID | rs34594498 | rs34410987 | rs33949390 | rs35801418 |
|---|---|---|---|---|
| PD_47 | C:C | C:C | G:G | A:A |
| PD_48 | C:C | C:C | G:G | A:A |
| PD_49 | C:C | C:C | G:G | A:A |
| PD_50 | C:C | C:C | G:G | A:A |
| PD_51 | C:C | C:C | G:G | A:A |
| PD_52 | C:C | C:C | G:G | A:A |
| PD_53 | C:C | C:C | G:G | A:A |
| PD_54 | C:C | C:C | G:G | A:A |
| PD_55 | C:C | C:C | G:G | A:A |
| PD_56 | C:C | C:C | G:G | A:A |
| PD_57 | C:C | C:C | G:G | A:A |
| PD_58 | C:C | C:C | G:G | A:A |
| PD_59 | C:C | C:C | G:G | A:A |
| PD_60 | C:C | C:C | G:G | A:A |
| PD_61 | C:C | C:C | G:G | A:A |
| PD_62 | C:C | C:C | G:G | A:A |
| PD_63 | C:C | C:C | G:G | A:A |
| PD_64 | C:C | C:C | G:G | A:A |
| PD_65 | C:C | C:C | G:G | A:A |
| PD_66 | C:C | C:C | G:G | A:A |
| PD_67 | C:C | C:C | G:G | A:A |
| PD_68 | C:C | C:C | G:G | A:A |
| PD_69 | C:C | C:C | G:G | A:A |
| PD_70 | C:C | C:C | G:G | A:A |
| PD_71 | C:C | C:C | G:G | A:A |
| PD_72 | C:C | C:C | G:G | A:A |
| PD_73 | C:C | C:C | G:G | A:A |
| PD_74 | C:C | C:C | G:G | A:A |
| PD_75 | C:C | C:C | G:G | A:A |
| PD_76 | C:C | C:C | G:G | A:A |
| PD_77 | C:C | C:C | G:G | A:A |
| PD_78 | C:C | C:C | G:G | A:A |
| PD_79 | C:C | C:C | G:G | A:A |
| PD_80 | C:C | C:C | G:G | A:A |
| PD_81 | C:C | C:C | G:G | A:A |
| PD_82 | C:C | C:C | G:G | A:A |
| PD_83 | C:C | C:C | G:G | A:A |
| PD_84 | C:C | C:C | G:G | A:A |
| PD_85 | C:C | C:C | G:G | A:A |
| PD_86 | C:C | C:C | G:G | A:A |
| PD_87 | C:C | C:C | G:G | A:A |
| PD_88 | C:C | C:C | G:G | A:A |
| PD_89 | C:C | C:C | G:G | A:A |
| PD_90 | C:C | C:C | G:G | A:A |
| PD_91 | C:C | C:C | G:G | A:A |
| PD_92 | C:C | C:C | G:G | A:A |

| Sample ID | rs34594498 | rs34410987 | rs33949390 | rs35801418 |
|---|---|---|---|---|
| Control 1 | C:C | C:C | G:G | A:A |
| Control 2 | C:C | C:C | G:G | A:A |
| Control 3 | C:C | C:C | G:G | A:A |
| Control 4 | C:C | C:C | G:G | A:A |
| Control 5 | C:C | C:C | G:G | A:A |
| Control 6 | C:C | C:C | G:G | A:A |
| Control 7 | C:C | C:C | G:G | A:A |
| Control 8 | C:C | C:C | G:G | A:A |
| Control 9 | C:C | C:C | G:G | A:A |
| Control 10 | C:C | C:C | G:G | A:A |
| Control 11 | C:C | C:C | G:G | A:A |
| Control 12 | C:C | C:C | G:G | A:A |
| Control 13 | C:C | C:C | G:G | A:A |
| Control 14 | C:C | C:C | G:G | A:A |
| Control 15 | C:C | C:C | G:G | A:A |
| Control 16 | C:C | C:C | G:G | A:A |
| Control 17 | C:C | C:C | G:G | A:A |
| Control 18 | C:C | C:C | G:G | A:A |
| Control 19 | C:C | C:C | G:G | A:A |
| Control 20 | C:C | C:C | G:G | A:A |
| Control 21 | C:C | C:C | G:G | A:A |
| Control 22 | C:C | C:C | G:G | A:A |
| Control 23 | C:C | C:C | G:G | A:A |
| Control 24 | C:C | C:C | G:G | A:A |
| Control 25 | C:C | C:C | G:G | A:A |
| Control 26 | C:C | C:C | G:G | A:A |
| Control 27 | C:C | C:C | G:G | A:A |
| Control 28 | C:C | C:C | G:G | A:A |
| Control 29 | C:C | C:C | G:G | A:A |
| Control 30 | C:C | C:C | G:G | A:A |
| Control 31 | C:C | C:C | G:G | A:A |
| Control 32 | C:C | C:C | G:G | A:A |
| Control 33 | C:C | C:C | G:G | A:A |
| Control 34 | C:C | C:C | G:G | A:A |
| Control 35 | C:C | C:C | G:G | A:A |
| Control 36 | C:C | C:C | G:G | A:A |
| Control 37 | C:C | C:C | G:G | A:A |
| Control 38 | C:C | C:C | G:G | A:A |
| Control 39 | C:C | C:C | G:G | A:A |
| Control 40 | C:C | C:C | G:G | A:A |
| Control 41 | C:C | C:C | G:G | A:A |
| Control 42 | C:C | C:C | G:G | A:A |
| Control 43 | C:C | C:C | G:G | A:A |
| Control 44 | C:C | C:C | G:G | A:A |
| Control 45 | C:C | C:C | G:G | A:A |
| Control 46 | C:C | C:C | G:G | A:A |

| Sample ID | rs34594498 | rs34410987 | rs33949390 | rs35801418 |
|---|---|---|---|---|
| Control 47 | C:C | C:C | G:G | A:A |
| Control 48 | C:C | C:C | G:G | A:A |
| Control 49 | C:C | C:C | G:G | A:A |
| Control 50 | C:C | C:C | G:G | A:A |
| Control 51 | C:C | C:C | G:G | A:A |
| Control 52 | C:C | C:C | G:G | A:A |
| Control 53 | C:C | C:C | G:G | A:A |
| Control 54 | C:C | C:C | G:G | A:A |
| Control 55 | C:C | C:C | G:G | A:A |
| Control 56 | C:C | C:C | G:G | A:A |
| Control 57 | C:C | C:C | G:G | A:A |
| Control 58 | C:C | C:C | G:G | A:A |
| Control 59 | C:C | C:C | G:G | A:A |
| Control 60 | C:C | C:C | G:G | A:A |
| Control 61 | C:C | C:C | G:G | A:A |
| Control 62 | C:C | C:C | G:G | A:A |
| Control 63 | C:C | C:C | G:G | A:A |
| Control 64 | C:C | C:C | G:G | A:A |
| Control 65 | C:C | C:C | G:G | A:A |
| Control 66 | C:C | C:C | G:G | A:A |
| Control 67 | C:C | C:C | G:G | A:A |
| Control 68 | C:C | C:C | G:G | A:A |
| Control 69 | C:C | C:C | G:G | A:A |
| Control 70 | C:C | C:C | G:G | A:A |
| Control 71 | C:C | C:C | G:G | A:A |
| Control 72 | C:C | C:C | G:G | A:A |
| Control 73 | C:C | C:C | G:G | A:A |
| Control 74 | C:C | C:C | G:G | A:A |
| Control 75 | C:C | C:C | G:G | A:A |
| Control 76 | C:C | C:C | G:G | A:A |
| Control 77 | C:C | C:C | G:G | A:A |
| Control 78 | C:C | C:C | G:G | A:A |
| Control 79 | C:C | C:C | G:G | A:A |
| Control 80 | C:C | C:C | G:G | A:A |
| Control 81 | C:C | C:C | G:G | A:A |
| Control 82 | C:C | C:C | G:G | A:A |
| Control 83 | C:C | C:C | G:G | A:A |
| Control 84 | C:C | C:C | G:G | A:A |
| Control 85 | C:C | C:C | G:G | A:A |
| Control 86 | C:C | C:C | G:G | A:A |
| Control 87 | C:C | C:C | G:G | A:A |
| Control 88 | C:C | C:C | G:G | A:A |
| Control 89 | C:C | C:C | G:G | A:A |
| Control 90 | C:C | C:C | G:G | A:A |
| Control 91 | C:C | C:C | G:G | A:A |
| Control 92 | C:C | C:C | G:G | A:A |

| Sample ID | rs34594498 | rs34410987 | rs33949390 | rs35801418 |
|---|---|---|---|---|
| Control 93 | C:C | C:C | G:G | A:A |
| Control 94 | C:C | C:C | G:G | A:A |
| Control 95 | C:C | C:C | G:G | A:A |
| Control 96 | C:C | C:C | G:G | A:A |
| Control 97 | C:C | C:C | G:G | A:A |
| Control 98 | C:C | C:C | G:G | A:A |
| Control 99 | C:C | C:C | G:G | A:A |
| Control 100 | C:C | C:C | G:G | A:A |
| Control 101 | C:C | C:C | G:G | A:A |
| Control 102 | C:C | C:C | G:G | A:A |
| Control 103 | C:C | C:C | G:G | A:A |
| Control 104 | C:C | C:C | G:G | A:A |
| Control 105 | C:C | C:C | G:G | A:A |
| Control 106 | C:C | C:C | G:G | A:A |
| Control 107 | C:C | C:C | G:G | A:A |
| Control 108 | C:C | C:C | G:G | A:A |
| Control 109 | C:C | C:C | G:G | A:A |
| Control 110 | C:C | C:C | G:G | A:A |
| Control 111 | C:C | C:C | G:G | A:A |
| Control 112 | C:C | C:C | G:G | A:A |
| Control 113 | C:C | C:C | G:G | A:A |
| Control 114 | C:C | C:C | G:G | A:A |
| Control 115 | C:C | C:C | G:G | A:A |
| Control 116 | C:C | C:C | G:G | A:A |
| Control 117 | C:C | C:C | G:G | A:A |
| Control 118 | C:C | C:C | G:G | A:A |
| Control 119 | C:C | C:C | G:G | A:A |
| Control 120 | C:C | C:C | G:G | A:A |
| Control 121 | C:C | C:C | G:G | A:A |
| Control 122 | C:C | C:C | G:G | A:A |
| Control 123 | C:C | C:C | G:G | A:A |
| Control 124 | C:C | C:C | G:G | A:A |
| Control 125 | C:C | C:C | G:G | A:A |
| Control 126 | C:C | C:C | G:G | A:A |
| Control 127 | C:C | C:C | G:G | A:A |
| Control 128 | C:C | C:C | G:G | A:A |
| Control 129 | C:C | C:C | G:G | A:A |
| Control 130 | C:C | C:C | G:G | A:A |
| Control 131 | C:C | C:C | G:G | A:A |
| Control 132 | C:C | C:C | G:G | A:A |
| Control 133 | C:C | C:C | G:G | A:A |
| Control 134 | C:C | C:C | G:G | A:A |
| Control 135 | C:C | C:C | G:G | A:A |
| Control 136 | C:C | C:C | G:G | A:A |
| Control 137 | C:C | C:C | G:G | A:A |
| Control 138 | C:C | C:C | G:G | A:A |

| Sample ID | rs34594498 | rs34410987 | rs33949390 | rs35801418 |
|---|---|---|---|---|
| Control 139 | C:C | C:C | G:G | A:A |
| Control 140 | C:C | C:C | G:G | A:A |
| Control 141 | C:C | C:C | G:G | A:A |
| Control 142 | C:C | C:C | G:G | A:A |
| Control 143 | C:C | C:C | G:G | A:A |
| Control 144 | C:C | C:C | G:G | A:A |
| Control 145 | C:C | C:C | G:G | A:A |
| Control 146 | C:C | C:C | G:G | A:A |
| Control 147 | C:C | C:C | G:G | A:A |
| Control 148 | C:C | C:C | G:G | A:A |
| Control 149 | C:C | C:C | G:G | A:A |
| Control 150 | C:C | C:C | G:G | A:A |
| Control 151 | C:C | C:C | G:G | A:A |
| Control 152 | C:C | C:C | G:G | A:A |
| Control 153 | C:C | C:C | G:G | A:A |
| Control 154 | C:C | C:C | G:G | A:A |
| Control 155 | C:C | C:C | G:G | A:A |
| Control 156 | C:C | C:C | G:G | A:A |
| Control 157 | C:C | C:C | G:G | A:A |
| Control 158 | C:C | C:C | G:G | A:A |
| Control 159 | C:C | C:C | G:G | A:A |
| Control 160 | C:C | C:C | G:G | A:A |
| Control 161 | C:C | C:C | G:G | A:A |
| Control 162 | C:C | C:C | G:G | A:A |
| Control 163 | C:C | C:C | G:G | A:A |
| Control 164 | C:C | C:C | G:G | A:A |
| Control 165 | C:C | C:C | G:G | A:A |
| Control 166 | C:C | C:C | G:G | A:A |
| Control 167 | C:C | C:C | G:G | A:A |
| Control 168 | C:C | C:C | G:G | A:A |
| Control 169 | C:C | C:C | G:G | A:A |
| Control 170 | C:C | C:C | G:G | A:A |
| Control 171 | C:C | C:C | G:G | A:A |
| Control 172 | C:C | C:C | G:G | A:A |
| Control 173 | C:C | C:C | G:G | A:A |
| Control 174 | C:C | C:C | Missing | A:A |
| Control 175 | C:C | C:C | G:G | A:A |
| Control 176 | C:C | C:C | G:G | A:A |
| Control 177 | C:C | C:C | G:G | A:A |
| Control 178 | C:C | C:C | G:G | A:A |
| Control 179 | C:C | C:C | G:G | A:A |
| Control 180 | C:C | C:C | G:G | A:A |
| Control 181 | C:C | C:C | G:G | A:A |
| Control 182 | C:C | C:C | G:G | A:A |
| Control 183 | C:C | C:C | G:G | A:A |
| Control 184 | C:C | C:C | G:G | A:A |

| Sample ID | rs34594498 | rs34410987 | rs33949390 | rs35801418 |
|---|---|---|---|---|
| Control 185 | C:C | C:C | G:G | A:A |
| Control 186 | C:C | C:C | G:G | A:A |
| Control 187 | C:C | C:C | G:G | A:A |
| Control 188 | C:C | C:C | G:G | A:A |
| Control 189 | C:C | C:C | G:G | A:A |
| Control 190 | C:C | C:C | G:G | A:A |
| Control 191 | C:C | C:C | G:G | A:A |
| Control 192 | C:C | C:C | G:G | A:A |
| Control 193 | C:C | C:C | G:G | A:A |
| Control 194 | C:C | C:C | G:G | A:A |
| Control 195 | C:C | C:C | G:G | A:A |
| Control 196 | C:C | C:C | G:G | A:A |
| Control 197 | C:C | C:C | G:G | A:A |
| Control 198 | C:C | C:C | G:G | A:A |
| Control 199 | C:C | C:C | G:G | A:A |
| Control 200 | C:C | C:C | G:G | A:A |
| Control 201 | C:C | C:C | G:G | A:A |
| Control 202 | C:C | C:C | G:G | A:A |
| Control 203 | C:C | C:C | G:G | A:A |
| Control 204 | C:C | C:C | G:G | A:A |
| Control 205 | C:C | C:C | G:G | A:A |
| Control 206 | C:C | C:C | G:G | A:A |
| Control 207 | C:C | C:C | G:G | A:A |
| Control 208 | C:C | C:C | G:G | A:A |
| Control 209 | C:C | C:C | G:G | A:A |
| Control 210 | C:C | C:C | G:G | A:A |

| Sample ID | rs34637584 | rs35870237 | rs34778348 | rs35808389 |
|---|---|---|---|---|
| PD_01 | G:G | T:T | G:G | A:A |
| PD_02 | G:G | T:T | G:G | A:A |
| PD_03 | G:G | T:T | G:G | A:A |
| PD_04 | G:G | T:T | G:G | A:A |
| PD_05 | G:G | T:T | G:G | A:A |
| PD_06 | G:G | T:T | G:G | A:A |
| PD_07 | G:G | T:T | G:G | A:A |
| PD_08 | G:G | T:T | G:G | A:A |
| PD_09 | G:G | T:T | G:G | A:A |
| PD_10 | G:G | T:T | G:G | A:A |
| PD_11 | G:G | T:T | G:G | A:A |
| PD_12 | G:G | T:T | G:G | A:A |
| PD_13 | G:G | T:T | G:G | A:A |
| PD_14 | G:G | T:T | G:G | A:A |
| PD_15 | G:G | T:T | G:G | A:A |
| PD_16 | G:G | T:T | G:G | A:A |
| PD_17 | G:G | T:T | G:G | A:A |
| PD_18 | G:G | T:T | G:G | A:A |
| PD_19 | G:G | T:T | G:G | A:A |
| PD_20 | G:G | T:T | G:G | A:A |
| PD_21 | G:G | T:T | G:G | A:A |
| PD_22 | G:G | T:T | G:G | A:A |
| PD_23 | G:G | T:T | G:G | A:A |
| PD_24 | G:G | T:T | G:G | A:A |
| PD_25 | G:G | T:T | G:G | A:A |
| PD_26 | G:G | T:T | G:G | A:A |
| PD_27 | G:G | T:T | G:G | A:A |
| PD_28 | G:G | T:T | G:G | A:A |
| PD_29 | G:G | T:T | G:G | A:A |
| PD_30 | G:G | T:T | G:G | A:A |
| PD_31 | G:G | T:T | G:G | A:A |
| PD_32 | G:G | T:T | G:G | A:A |
| PD_33 | G:G | T:T | G:G | A:A |
| PD_34 | G:G | T:T | G:G | A:A |
| PD_35 | G:G | T:T | G:G | A:A |
| PD_36 | G:G | T:T | G:G | A:A |
| PD_37 | G:G | T:T | G:G | A:A |
| PD_38 | G:G | T:T | G:G | A:A |
| PD_39 | G:G | T:T | G:G | A:A |
| PD_40 | G:G | T:T | G:G | A:A |
| PD_41 | G:G | T:T | G:G | A:A |
| PD_42 | G:G | T:T | G:G | A:A |
| PD_43 | G:G | T:T | G:G | A:A |
| PD_44 | G:G | T:T | G:G | A:A |
| PD_45 | G:G | T:T | G:G | A:A |
| PD_46 | G:G | T:T | G:G | A:A |

| Sample ID | rs34637584 | rs35870237 | rs34778348 | rs35808389 |
|---|---|---|---|---|
| PD_47 | G:G | T:T | G:G | A:A |
| PD_48 | G:G | T:T | G:G | A:A |
| PD_49 | G:G | T:T | G:G | A:A |
| PD_50 | G:G | T:T | G:G | A:A |
| PD_51 | G:G | T:T | G:G | A:A |
| PD_52 | G:G | T:T | G:G | A:A |
| PD_53 | G:G | T:T | G:G | A:A |
| PD_54 | G:G | T:T | G:G | A:A |
| PD_55 | G:G | T:T | G:G | A:A |
| PD_56 | G:G | T:T | G:G | A:A |
| PD_57 | G:G | T:T | G:G | A:A |
| PD_58 | G:G | T:T | G:G | A:A |
| PD_59 | G:G | T:T | G:G | A:A |
| PD_60 | G:G | T:T | G:G | A:A |
| PD_61 | G:G | T:T | G:G | A:A |
| PD_62 | G:G | T:T | G:G | A:A |
| PD_63 | G:G | T:T | G:G | A:A |
| PD_64 | G:G | T:T | G:G | A:A |
| PD_65 | G:G | T:T | G:G | A:A |
| PD_66 | G:G | T:T | G:G | A:A |
| PD_67 | G:G | T:T | G:G | A:A |
| PD_68 | G:G | T:T | G:G | A:A |
| PD_69 | G:G | T:T | G:G | A:A |
| PD_70 | G:G | T:T | G:G | A:A |
| PD_71 | G:G | T:T | G:G | A:A |
| PD_72 | G:G | T:T | G:G | A:A |
| PD_73 | G:G | T:T | G:G | A:A |
| PD_74 | G:G | T:T | G:G | A:A |
| PD_75 | G:G | T:T | G:G | A:A |
| PD_76 | G:G | T:T | G:G | A:A |
| PD_77 | G:G | T:T | G:G | A:A |
| PD_78 | G:G | T:T | G:G | A:A |
| PD_79 | G:G | T:T | G:G | A:A |
| PD_80 | G:G | T:T | G:G | A:A |
| PD_81 | G:G | T:T | G:G | A:A |
| PD_82 | G:G | T:T | G:G | A:A |
| PD_83 | G:G | T:T | G:G | A:A |
| PD_84 | G:G | T:T | G:G | A:A |
| PD_85 | G:G | T:T | G:G | A:A |
| PD_86 | G:G | T:T | G:G | A:A |
| PD_87 | G:G | T:T | G:G | A:A |
| PD_88 | G:G | T:T | G:G | A:A |
| PD_89 | G:G | T:T | G:G | A:A |
| PD_90 | G:G | T:T | G:G | A:A |
| PD_91 | G:G | T:T | G:G | A:A |
| PD_92 | G:G | T:T | G:G | A:A |

| **Sample ID** | **rs34637584** | **rs35870237** | **rs34778348** | **rs35808389** |
|---|---|---|---|---|
| Control 1 | G:G | T:T | G:G | A:A |
| Control 2 | G:G | T:T | G:G | A:A |
| Control 3 | G:G | T:T | G:G | A:A |
| Control 4 | G:G | T:T | G:G | A:A |
| Control 5 | G:G | T:T | G:G | A:A |
| Control 6 | G:G | T:T | G:G | A:A |
| Control 7 | G:G | T:T | G:G | A:A |
| Control 8 | G:G | T:T | G:G | A:A |
| Control 9 | G:G | T:T | Missing | A:A |
| Control 10 | G:G | T:T | G:G | A:A |
| Control 11 | G:G | T:T | G:G | A:A |
| Control 12 | G:G | T:T | G:G | A:A |
| Control 13 | G:G | T:T | G:G | A:A |
| Control 14 | G:G | T:T | G:G | A:A |
| Control 15 | G:G | T:T | G:G | A:A |
| Control 16 | G:G | T:T | G:G | A:A |
| Control 17 | G:G | T:T | G:G | A:A |
| Control 18 | G:G | T:T | G:G | A:A |
| Control 19 | G:G | T:T | G:G | A:A |
| Control 20 | G:G | T:T | G:G | A:A |
| Control 21 | G:G | T:T | G:G | A:A |
| Control 22 | G:G | T:T | G:G | A:A |
| Control 23 | G:G | T:T | G:G | A:A |
| Control 24 | G:G | T:T | G:G | A:A |
| Control 25 | G:G | T:T | G:G | A:A |
| Control 26 | G:G | T:T | G:G | A:A |
| Control 27 | G:G | T:T | G:G | A:A |
| Control 28 | G:G | T:T | G:G | A:A |
| Control 29 | G:G | T:T | G:G | A:A |
| Control 30 | G:G | T:T | G:G | A:A |
| Control 31 | G:G | T:T | G:G | A:A |
| Control 32 | G:G | T:T | G:G | A:A |
| Control 33 | G:G | T:T | G:G | A:A |
| Control 34 | G:G | T:T | G:G | A:A |
| Control 35 | G:G | T:T | G:G | A:A |
| Control 36 | G:G | T:T | G:G | A:A |
| Control 37 | G:G | T:T | G:G | A:A |
| Control 38 | G:G | T:T | G:G | A:A |
| Control 39 | G:G | T:T | G:G | A:A |
| Control 40 | G:G | T:T | G:G | A:A |
| Control 41 | G:G | T:T | G:G | A:A |
| Control 42 | G:G | T:T | G:G | A:A |
| Control 43 | G:G | T:T | G:G | A:A |
| Control 44 | G:G | T:T | G:G | A:A |
| Control 45 | G:G | T:T | G:G | A:A |
| Control 46 | G:G | T:T | G:G | A:A |

| Sample ID | rs34637584 | rs35870237 | rs34778348 | rs35808389 |
|---|---|---|---|---|
| Control 47 | G:G | T:T | G:G | A:A |
| Control 48 | G:G | T:T | G:G | A:A |
| Control 49 | G:G | T:T | G:G | A:A |
| Control 50 | G:G | T:T | G:G | A:A |
| Control 51 | G:G | T:T | G:G | A:A |
| Control 52 | G:G | T:T | G:G | A:A |
| Control 53 | G:G | T:T | G:G | A:A |
| Control 54 | G:G | T:T | G:G | A:A |
| Control 55 | G:G | T:T | G:G | A:A |
| Control 56 | G:G | T:T | G:G | A:A |
| Control 57 | G:G | T:T | G:G | A:A |
| Control 58 | G:G | T:T | G:G | A:A |
| Control 59 | G:G | T:T | G:G | A:A |
| Control 60 | G:G | T:T | G:G | A:A |
| Control 61 | G:G | T:T | G:G | A:A |
| Control 62 | G:G | T:T | G:G | A:A |
| Control 63 | G:G | T:T | G:G | A:A |
| Control 64 | G:G | T:T | G:G | A:A |
| Control 65 | G:G | T:T | G:G | A:A |
| Control 66 | G:G | T:T | G:G | A:A |
| Control 67 | G:G | T:T | G:G | A:A |
| Control 68 | G:G | T:T | G:G | A:A |
| Control 69 | G:G | T:T | G:G | A:A |
| Control 70 | G:G | T:T | G:G | A:A |
| Control 71 | G:G | T:T | G:G | A:A |
| Control 72 | G:G | T:T | G:G | A:A |
| Control 73 | G:G | T:T | G:G | A:A |
| Control 74 | G:G | T:T | G:G | A:A |
| Control 75 | G:G | T:T | G:G | A:A |
| Control 76 | G:G | T:T | G:G | A:A |
| Control 77 | G:G | T:T | G:G | A:A |
| Control 78 | G:G | T:T | G:G | A:A |
| Control 79 | G:G | T:T | G:G | A:A |
| Control 80 | G:G | T:T | G:G | A:A |
| Control 81 | G:G | T:T | G:G | A:A |
| Control 82 | G:G | T:T | G:G | A:A |
| Control 83 | G:G | T:T | G:G | A:A |
| Control 84 | G:G | T:T | G:G | A:A |
| Control 85 | G:G | T:T | G:G | A:A |
| Control 86 | G:G | T:T | G:G | A:A |
| Control 87 | G:G | T:T | G:G | A:A |
| Control 88 | G:G | T:T | G:G | A:A |
| Control 89 | G:G | T:T | G:G | A:A |
| Control 90 | G:G | T:T | G:G | A:A |
| Control 91 | G:G | T:T | G:G | A:A |
| Control 92 | G:G | T:T | G:G | A:A |

| Sample ID | rs34637584 | rs35870237 | rs34778348 | rs35808389 |
|---|---|---|---|---|
| Control 93 | G:G | T:T | G:G | A:A |
| Control 94 | G:G | T:T | G:G | A:A |
| Control 95 | G:G | T:T | G:G | A:A |
| Control 96 | G:G | T:T | G:G | A:A |
| Control 97 | G:G | T:T | G:G | A:A |
| Control 98 | G:G | T:T | G:G | A:A |
| Control 99 | G:G | Missing | G:G | A:A |
| Control 100 | G:G | T:T | G:G | A:A |
| Control 101 | G:G | T:T | G:G | A:A |
| Control 102 | G:G | T:T | G:G | A:A |
| Control 103 | G:G | T:T | G:G | A:A |
| Control 104 | G:G | T:T | G:G | A:A |
| Control 105 | G:G | T:T | G:G | A:A |
| Control 106 | G:G | T:T | G:G | A:A |
| Control 107 | G:G | T:T | G:G | A:A |
| Control 108 | G:G | T:T | G:G | A:A |
| Control 109 | G:G | T:T | G:G | A:A |
| Control 110 | G:G | T:T | G:G | A:A |
| Control 111 | G:G | T:T | G:G | A:A |
| Control 112 | G:G | T:T | G:G | A:A |
| Control 113 | G:G | T:T | G:G | A:A |
| Control 114 | G:G | T:T | G:G | A:A |
| Control 115 | G:G | T:T | G:G | A:A |
| Control 116 | G:G | T:T | G:G | A:A |
| Control 117 | G:G | Missing | G:G | A:A |
| Control 118 | G:G | T:T | G:G | A:A |
| Control 119 | G:G | T:T | G:G | A:A |
| Control 120 | G:G | T:T | G:G | A:A |
| Control 121 | G:G | T:T | G:G | A:A |
| Control 122 | G:G | T:T | G:G | A:A |
| Control 123 | G:G | T:T | G:G | A:A |
| Control 124 | G:G | T:T | G:G | A:A |
| Control 125 | G:G | T:T | G:G | A:A |
| Control 126 | G:G | T:T | G:G | A:A |
| Control 127 | G:G | T:T | G:G | A:A |
| Control 128 | G:G | T:T | G:G | A:A |
| Control 129 | G:G | T:T | G:G | A:A |
| Control 130 | G:G | T:T | G:G | A:A |
| Control 131 | G:G | T:T | G:G | A:A |
| Control 132 | G:G | T:T | G:G | A:A |
| Control 133 | G:G | T:T | G:G | A:A |
| Control 134 | G:G | T:T | G:G | A:A |
| Control 135 | G:G | T:T | G:G | A:A |
| Control 136 | G:G | T:T | G:G | A:A |
| Control 137 | G:G | T:T | G:G | A:A |
| Control 138 | G:G | T:T | G:G | A:A |

| Sample ID | rs34637584 | rs35870237 | rs34778348 | rs35808389 |
|---|---|---|---|---|
| Control 139 | G:G | T:T | G:G | A:A |
| Control 140 | G:G | T:T | G:G | A:A |
| Control 141 | G:G | T:T | G:G | A:A |
| Control 142 | G:G | T:T | G:G | A:A |
| Control 143 | G:G | T:T | G:G | A:A |
| Control 144 | G:G | T:T | G:G | A:A |
| Control 145 | G:G | T:T | G:G | A:A |
| Control 146 | G:G | T:T | G:G | A:A |
| Control 147 | G:G | T:T | G:G | A:A |
| Control 148 | G:G | T:T | G:G | A:A |
| Control 149 | G:G | T:T | G:G | A:A |
| Control 150 | G:G | T:T | G:G | A:A |
| Control 151 | G:G | T:T | G:G | A:A |
| Control 152 | G:G | T:T | G:G | A:A |
| Control 153 | G:G | T:T | G:G | A:A |
| Control 154 | G:G | T:T | G:G | A:A |
| Control 155 | G:G | T:T | G:G | A:A |
| Control 156 | G:G | T:T | G:G | A:A |
| Control 157 | G:G | T:T | G:G | A:A |
| Control 158 | G:G | T:T | G:G | A:A |
| Control 159 | G:G | T:T | G:G | A:A |
| Control 160 | G:G | T:T | G:G | A:A |
| Control 161 | G:G | T:T | G:G | A:A |
| Control 162 | G:G | T:T | G:G | A:A |
| Control 163 | G:G | T:T | G:G | A:A |
| Control 164 | G:G | T:T | G:G | A:A |
| Control 165 | G:G | T:T | G:G | A:A |
| Control 166 | G:G | T:T | G:G | A:A |
| Control 167 | G:G | T:T | G:G | A:A |
| Control 168 | G:G | T:T | G:G | A:A |
| Control 169 | G:G | T:T | G:G | A:A |
| Control 170 | G:G | T:T | G:G | A:A |
| Control 171 | G:G | T:T | G:G | A:A |
| Control 172 | G:G | T:T | G:G | A:A |
| Control 173 | G:G | T:T | G:G | A:A |
| Control 174 | G:G | T:T | G:G | A:A |
| Control 175 | G:G | T:T | G:G | A:A |
| Control 176 | G:G | T:T | G:G | A:A |
| Control 177 | G:G | T:T | G:G | A:A |
| Control 178 | G:G | T:T | G:G | A:A |
| Control 179 | G:G | T:T | G:G | A:A |
| Control 180 | G:G | T:T | G:G | A:A |
| Control 181 | G:G | T:T | G:G | A:A |
| Control 182 | G:G | T:T | G:G | A:A |
| Control 183 | G:G | T:T | G:G | A:A |
| Control 184 | G:G | T:T | G:G | A:A |

| **Sample ID** | **rs34637584** | **rs35870237** | **rs34778348** | **rs35808389** |
|---|---|---|---|---|
| Control 185 | G:G | T:T | G:G | A:A |
| Control 186 | G:G | T:T | G:G | A:A |
| Control 187 | G:G | T:T | G:G | A:A |
| Control 188 | G:G | T:T | G:G | A:A |
| Control 189 | G:G | T:T | G:G | A:A |
| Control 190 | G:G | T:T | G:G | A:A |
| Control 191 | G:G | T:T | G:G | A:A |
| Control 192 | G:G | T:T | G:G | A:A |
| Control 193 | G:G | T:T | G:G | A:A |
| Control 194 | G:G | T:T | G:G | A:A |
| Control 195 | G:G | T:T | G:G | A:A |
| Control 196 | G:G | T:T | G:G | A:A |
| Control 197 | G:G | T:T | G:G | A:A |
| Control 198 | G:G | T:T | G:G | A:A |
| Control 199 | G:G | T:T | G:G | A:A |
| Control 200 | G:G | T:T | G:G | A:A |
| Control 201 | G:G | T:T | G:G | A:A |
| Control 202 | G:G | T:T | G:G | A:A |
| Control 203 | G:G | T:T | G:G | A:A |
| Control 204 | G:G | T:T | G:G | A:A |
| Control 205 | G:G | T:T | G:G | A:A |
| Control 206 | G:G | T:T | G:G | A:A |
| Control 207 | G:G | T:T | G:G | A:A |
| Control 208 | G:G | T:T | G:G | A:A |
| Control 209 | G:G | T:T | G:G | A:A |
| Control 210 | G:G | T:T | G:G | A:A |

| Sample ID | rs34805604 | rs74163686 | rs34995376 | rs281865052 |
|---|---|---|---|---|
| PD_01 | A:A | A:A | G:G | A:A |
| PD_02 | A:A | A:A | G:G | A:A |
| PD_03 | A:A | A:A | G:G | A:A |
| PD_04 | A:A | A:A | G:G | A:A |
| PD_05 | A:A | A:A | G:G | A:A |
| PD_06 | A:A | A:A | G:G | A:A |
| PD_07 | A:A | A:A | G:G | A:A |
| PD_08 | A:A | A:A | G:G | A:A |
| PD_09 | A:A | A:A | G:G | A:A |
| PD_10 | A:A | A:A | G:G | A:A |
| PD_11 | A:A | A:A | G:G | A:A |
| PD_12 | A:A | A:A | G:G | A:A |
| PD_13 | A:A | A:A | G:G | A:A |
| PD_14 | A:A | A:A | G:G | A:A |
| PD_15 | A:A | A:A | G:G | A:A |
| PD_16 | A:A | A:A | G:G | A:A |
| PD_17 | A:A | A:A | G:G | A:A |
| PD_18 | A:A | A:A | G:G | A:A |
| PD_19 | A:A | A:A | G:G | A:A |
| PD_20 | A:A | A:A | G:G | A:A |
| PD_21 | A:A | A:A | G:G | A:A |
| PD_22 | A:A | A:A | G:G | A:A |
| PD_23 | A:A | A:A | G:G | A:A |
| PD_24 | A:A | A:A | G:G | A:A |
| PD_25 | A:A | A:A | G:G | A:A |
| PD_26 | A:A | A:A | G:G | A:A |
| PD_27 | A:A | A:A | G:G | A:A |
| PD_28 | A:A | A:A | G:G | A:A |
| PD_29 | A:A | A:A | G:G | A:A |
| PD_30 | A:A | A:A | G:G | A:A |
| PD_31 | A:A | A:A | G:G | A:A |
| PD_32 | A:A | A:A | G:G | A:A |
| PD_33 | A:A | A:A | G:G | A:A |
| PD_34 | A:A | A:A | G:G | A:A |
| PD_35 | A:A | A:A | G:G | A:A |
| PD_36 | A:A | A:A | G:G | A:A |
| PD_37 | A:A | A:A | G:G | A:A |
| PD_38 | A:A | A:A | G:G | A:A |
| PD_39 | A:A | A:A | G:G | A:A |
| PD_40 | A:A | A:A | G:G | A:A |
| PD_41 | A:A | A:A | G:G | A:A |
| PD_42 | A:A | A:A | G:G | A:A |
| PD_43 | A:A | A:A | G:G | A:A |
| PD_44 | A:A | A:A | G:G | A:A |
| PD_45 | A:A | A:A | G:G | A:A |
| PD_46 | A:A | A:A | G:G | A:A |

| Sample ID | rs34805604 | rs74163686 | rs34995376 | rs281865052 |
|---|---|---|---|---|
| PD_47 | A:A | A:A | G:G | A:A |
| PD_48 | A:A | A:A | G:G | A:A |
| PD_49 | A:A | A:A | G:G | A:A |
| PD_50 | A:A | A:A | G:G | A:A |
| PD_51 | A:A | A:A | G:G | A:A |
| PD_52 | A:A | A:A | G:G | A:A |
| PD_53 | A:A | A:A | G:G | A:A |
| PD_54 | A:A | A:A | G:G | A:A |
| PD_55 | A:A | A:A | G:G | A:A |
| PD_56 | A:A | A:A | G:G | A:A |
| PD_57 | A:A | A:A | G:G | A:A |
| PD_58 | A:A | A:A | G:G | A:A |
| PD_59 | A:A | A:A | G:G | A:A |
| PD_60 | A:A | A:A | G:G | A:A |
| PD_61 | A:A | A:A | G:G | A:A |
| PD_62 | A:A | A:A | G:G | A:A |
| PD_63 | A:A | A:A | G:G | A:A |
| PD_64 | A:A | A:A | G:G | A:A |
| PD_65 | A:A | A:A | G:G | A:A |
| PD_66 | A:A | A:A | G:G | A:A |
| PD_67 | A:A | A:A | G:G | A:A |
| PD_68 | A:A | A:A | G:G | A:A |
| PD_69 | A:A | A:A | G:G | A:A |
| PD_70 | A:A | A:A | G:G | A:A |
| PD_71 | A:A | A:A | G:G | A:A |
| PD_72 | A:A | A:A | G:G | A:A |
| PD_73 | A:A | A:A | G:G | A:A |
| PD_74 | A:A | A:A | G:G | A:A |
| PD_75 | A:A | A:A | G:G | A:A |
| PD_76 | A:A | A:A | G:G | A:A |
| PD_77 | A:A | A:A | G:G | A:A |
| PD_78 | A:A | A:A | G:G | A:A |
| PD_79 | A:A | A:A | G:G | A:A |
| PD_80 | A:A | A:A | G:G | A:A |
| PD_81 | A:A | A:A | G:G | A:A |
| PD_82 | A:A | A:A | G:G | A:A |
| PD_83 | A:A | A:A | G:G | A:A |
| PD_84 | A:A | A:A | G:G | A:A |
| PD_85 | A:A | A:A | G:G | A:A |
| PD_86 | A:A | A:A | G:G | A:A |
| PD_87 | A:A | A:A | G:G | A:A |
| PD_88 | A:A | A:A | G:G | A:A |
| PD_89 | A:A | A:A | G:G | A:A |
| PD_90 | A:A | A:A | G:G | A:A |
| PD_91 | A:A | A:A | G:G | A:A |
| PD_92 | A:A | A:A | G:G | A:A |

| Sample ID | rs34805604 | rs74163686 | rs34995376 | rs281865052 |
|---|---|---|---|---|
| Control 1 | A:A | A:A | G:G | A:A |
| Control 2 | A:A | A:A | G:G | A:A |
| Control 3 | A:A | A:A | G:G | A:A |
| Control 4 | A:A | A:A | G:G | A:A |
| Control 5 | A:A | A:A | G:G | A:A |
| Control 6 | A:A | A:A | G:G | A:A |
| Control 7 | A:A | A:A | G:G | A:A |
| Control 8 | A:A | A:A | G:G | A:A |
| Control 9 | A:A | A:A | G:G | A:A |
| Control 10 | A:A | A:A | G:G | A:A |
| Control 11 | A:A | A:A | G:G | A:A |
| Control 12 | A:A | A:A | G:G | A:A |
| Control 13 | A:A | A:A | G:G | A:A |
| Control 14 | A:A | A:A | G:G | A:A |
| Control 15 | A:A | A:A | G:G | A:A |
| Control 16 | A:A | A:A | G:G | A:A |
| Control 17 | A:A | A:A | G:G | A:A |
| Control 18 | A:A | A:A | G:G | A:A |
| Control 19 | A:A | A:A | G:G | A:A |
| Control 20 | A:A | A:A | G:G | A:A |
| Control 21 | A:A | A:A | G:G | A:A |
| Control 22 | A:A | A:A | G:G | A:A |
| Control 23 | A:A | A:A | G:G | A:A |
| Control 24 | A:A | A:A | G:G | A:A |
| Control 25 | A:A | A:A | G:G | A:A |
| Control 26 | A:A | A:A | G:G | A:A |
| Control 27 | A:A | A:A | G:G | A:A |
| Control 28 | A:A | A:A | G:G | A:A |
| Control 29 | A:A | A:A | G:G | A:A |
| Control 30 | A:A | A:A | G:G | A:A |
| Control 31 | A:A | A:A | G:G | A:A |
| Control 32 | A:A | A:A | G:G | A:A |
| Control 33 | A:A | A:A | G:G | A:A |
| Control 34 | A:A | A:A | G:G | A:A |
| Control 35 | A:A | A:A | G:G | A:A |
| Control 36 | A:A | A:A | G:G | A:A |
| Control 37 | A:A | A:A | G:G | A:A |
| Control 38 | A:A | A:A | G:G | A:A |
| Control 39 | A:A | A:A | G:G | A:A |
| Control 40 | A:A | A:A | G:G | A:A |
| Control 41 | A:A | A:A | G:G | A:A |
| Control 42 | A:A | A:A | G:G | A:A |
| Control 43 | A:A | A:A | G:G | A:A |
| Control 44 | A:A | A:A | G:G | A:A |
| Control 45 | A:A | A:A | G:G | A:A |
| Control 46 | A:A | A:A | G:G | A:A |

| Sample ID | rs34805604 | rs74163686 | rs34995376 | rs281865052 |
|---|---|---|---|---|
| Control 47 | A:A | A:A | G:G | A:A |
| Control 48 | A:A | A:A | G:G | A:A |
| Control 49 | A:A | A:A | G:G | A:A |
| Control 50 | A:A | A:A | G:G | A:A |
| Control 51 | A:A | A:A | G:G | A:A |
| Control 52 | A:A | A:A | G:G | A:A |
| Control 53 | A:A | A:A | G:G | A:A |
| Control 54 | A:A | A:A | G:G | A:A |
| Control 55 | A:A | A:A | G:G | A:A |
| Control 56 | A:A | A:A | G:G | A:A |
| Control 57 | A:A | A:A | G:G | A:A |
| Control 58 | A:A | A:A | G:G | A:A |
| Control 59 | A:A | A:A | G:G | A:A |
| Control 60 | A:A | A:A | G:G | A:A |
| Control 61 | A:A | A:A | G:G | A:A |
| Control 62 | A:A | A:A | G:G | A:A |
| Control 63 | A:A | A:A | G:G | A:A |
| Control 64 | A:A | A:A | G:G | A:A |
| Control 65 | A:A | A:A | G:G | A:A |
| Control 66 | A:A | A:A | G:G | A:A |
| Control 67 | A:A | A:A | G:G | A:A |
| Control 68 | A:A | A:A | G:G | A:A |
| Control 69 | A:A | A:A | G:G | A:A |
| Control 70 | A:A | A:A | G:G | A:A |
| Control 71 | A:A | A:A | G:G | A:A |
| Control 72 | A:A | A:A | G:G | A:A |
| Control 73 | A:A | A:A | G:G | A:A |
| Control 74 | A:A | A:A | G:G | A:A |
| Control 75 | A:A | A:A | G:G | A:A |
| Control 76 | A:A | A:A | G:G | A:A |
| Control 77 | A:A | A:A | G:G | A:A |
| Control 78 | A:A | A:A | G:G | A:A |
| Control 79 | A:A | A:A | G:G | A:A |
| Control 80 | A:A | A:A | G:G | A:A |
| Control 81 | A:A | A:A | G:G | A:A |
| Control 82 | A:A | A:A | G:G | A:A |
| Control 83 | A:A | A:A | G:G | A:A |
| Control 84 | A:A | A:A | G:G | A:A |
| Control 85 | A:A | A:A | G:G | A:A |
| Control 86 | A:A | A:A | G:G | A:A |
| Control 87 | A:A | A:A | G:G | A:A |
| Control 88 | A:A | A:A | G:G | A:A |
| Control 89 | A:A | A:A | G:G | A:A |
| Control 90 | A:A | A:A | G:G | A:A |
| Control 91 | A:A | A:A | G:G | A:A |
| Control 92 | A:A | A:A | G:G | A:A |

| Sample ID | rs34805604 | rs74163686 | rs34995376 | rs281865052 |
|---|---|---|---|---|
| Control 93 | A:A | A:A | G:G | A:A |
| Control 94 | A:A | A:A | G:G | A:A |
| Control 95 | A:A | A:A | G:G | A:A |
| Control 96 | A:A | A:A | G:G | A:A |
| Control 97 | A:A | A:A | G:G | A:A |
| Control 98 | A:A | A:A | G:G | A:A |
| Control 99 | A:A | A:A | G:G | A:A |
| Control 100 | A:A | A:A | G:G | A:A |
| Control 101 | A:A | A:A | G:G | A:A |
| Control 102 | A:A | A:A | G:G | A:A |
| Control 103 | A:A | A:A | G:G | A:A |
| Control 104 | A:A | A:A | G:G | A:A |
| Control 105 | A:A | A:A | G:G | A:A |
| Control 106 | A:A | A:A | G:G | A:A |
| Control 107 | A:A | A:A | G:G | A:A |
| Control 108 | A:A | A:A | G:G | A:A |
| Control 109 | A:A | A:A | G:G | A:A |
| Control 110 | A:A | A:A | G:G | A:A |
| Control 111 | A:A | A:A | G:G | A:A |
| Control 112 | A:A | A:A | G:G | A:A |
| Control 113 | A:A | A:A | G:G | A:A |
| Control 114 | A:A | A:A | G:G | A:A |
| Control 115 | A:A | A:A | G:G | A:A |
| Control 116 | A:A | A:A | G:G | A:A |
| Control 117 | A:A | A:A | G:G | A:A |
| Control 118 | A:A | A:A | G:G | A:A |
| Control 119 | A:A | A:A | G:G | A:A |
| Control 120 | A:A | A:A | G:G | A:A |
| Control 121 | A:A | A:A | G:G | A:A |
| Control 122 | A:A | A:A | G:G | A:A |
| Control 123 | A:A | A:A | G:G | A:A |
| Control 124 | A:A | A:A | G:G | A:A |
| Control 125 | A:A | A:A | G:G | A:A |
| Control 126 | A:A | A:A | G:G | A:A |
| Control 127 | A:A | A:A | G:G | A:A |
| Control 128 | A:A | A:A | G:G | A:A |
| Control 129 | A:A | A:A | G:G | A:A |
| Control 130 | A:A | A:A | G:G | A:A |
| Control 131 | A:A | A:A | G:G | A:A |
| Control 132 | A:A | A:A | G:G | A:A |
| Control 133 | A:A | A:A | G:G | A:A |
| Control 134 | A:A | A:A | G:G | A:A |
| Control 135 | A:A | A:A | G:G | A:A |
| Control 136 | A:A | A:A | G:G | A:A |
| Control 137 | A:A | A:A | G:G | A:A |
| Control 138 | A:A | A:A | G:G | A:A |

| Sample ID | rs34805604 | rs74163686 | rs34995376 | rs281865052 |
|---|---|---|---|---|
| Control 139 | A:A | A:A | G:G | A:A |
| Control 140 | A:A | A:A | G:G | A:A |
| Control 141 | A:A | A:A | G:G | A:A |
| Control 142 | A:A | A:A | G:G | A:A |
| Control 143 | A:A | A:A | G:G | A:A |
| Control 144 | A:A | A:A | G:G | A:A |
| Control 145 | A:A | A:A | G:G | A:A |
| Control 146 | A:A | A:A | G:G | A:A |
| Control 147 | A:A | A:A | G:G | A:A |
| Control 148 | A:A | A:A | G:G | A:A |
| Control 149 | A:A | A:A | G:G | A:A |
| Control 150 | A:A | A:A | G:G | A:A |
| Control 151 | A:A | A:A | G:G | A:A |
| Control 152 | A:A | A:A | G:G | A:A |
| Control 153 | A:A | A:A | G:G | A:A |
| Control 154 | A:A | A:A | G:G | A:A |
| Control 155 | A:A | A:A | G:G | Missing |
| Control 156 | A:A | A:A | G:G | A:A |
| Control 157 | A:A | A:A | G:G | A:A |
| Control 158 | A:A | A:A | G:G | A:A |
| Control 159 | A:A | A:A | G:G | A:A |
| Control 160 | A:A | A:A | G:G | A:A |
| Control 161 | A:A | A:A | G:G | A:A |
| Control 162 | Missing | A:A | G:G | A:A |
| Control 163 | A:A | A:A | G:G | A:A |
| Control 164 | A:A | A:A | G:G | A:A |
| Control 165 | A:A | A:A | G:G | A:A |
| Control 166 | A:A | A:A | G:G | A:A |
| Control 167 | A:A | A:A | G:G | Missing |
| Control 168 | A:A | A:A | G:G | A:A |
| Control 169 | A:A | A:A | G:G | A:A |
| Control 170 | A:A | A:A | G:G | A:A |
| Control 171 | A:A | A:A | G:G | A:A |
| Control 172 | A:A | A:A | G:G | A:A |
| Control 173 | A:A | A:A | G:G | A:A |
| Control 174 | A:A | A:A | G:G | A:A |
| Control 175 | A:A | A:A | G:G | A:A |
| Control 176 | A:A | A:A | G:G | A:A |
| Control 177 | A:A | A:A | G:G | A:A |
| Control 178 | A:A | A:A | G:G | A:A |
| Control 179 | A:A | A:A | G:G | A:A |
| Control 180 | A:A | A:A | G:G | A:A |
| Control 181 | A:A | A:A | G:G | A:A |
| Control 182 | A:A | A:A | G:G | A:A |
| Control 183 | A:A | A:A | G:G | A:A |
| Control 184 | A:A | A:A | G:G | A:A |

| Sample ID | rs34805604 | rs74163686 | rs34995376 | rs281865052 |
|---|---|---|---|---|
| Control 185 | A:A | A:A | G:G | A:A |
| Control 186 | A:A | A:A | G:G | A:A |
| Control 187 | A:A | A:A | G:G | A:A |
| Control 188 | A:A | A:A | G:G | A:A |
| Control 189 | A:A | A:A | G:G | A:A |
| Control 190 | A:A | A:A | G:G | A:A |
| Control 191 | A:A | A:A | G:G | A:A |
| Control 192 | A:A | A:A | G:G | A:A |
| Control 193 | A:A | A:A | G:G | A:A |
| Control 194 | A:A | A:A | G:G | A:A |
| Control 195 | A:A | A:A | G:G | A:A |
| Control 196 | A:A | A:A | G:G | A:A |
| Control 197 | A:A | A:A | G:G | A:A |
| Control 198 | A:A | A:A | G:G | A:A |
| Control 199 | A:A | A:A | G:G | A:A |
| Control 200 | Missing | A:A | G:G | A:A |
| Control 201 | A:A | A:A | G:G | A:A |
| Control 202 | A:A | A:A | G:G | A:A |
| Control 203 | A:A | A:A | G:G | A:A |
| Control 204 | A:A | A:A | G:G | A:A |
| Control 205 | A:A | A:A | G:G | A:A |
| Control 206 | A:A | A:A | G:G | A:A |
| Control 207 | A:A | A:A | G:G | A:A |
| Control 208 | A:A | A:A | G:G | A:A |
| Control 209 | A:A | A:A | G:G | A:A |
| Control 210 | A:A | A:A | G:G | A:A |
